# Antagonist tendon vibration dampens estimates of persistent inward currents in motor units of the human lower limb

**DOI:** 10.1101/2022.08.02.502526

**Authors:** Gregory E P Pearcey, Obaid U Khurram, James A Beauchamp, Francesco Negro, Charles J Heckman

**Author notes:** **Address for correspondence:** Gregory Pearcey, School of Human Kinetics and Recreation, Memorial University of Newfoundland, St. John’s, NL, Canada, A1C 5S7.

## Abstract

We can readily measure motoneuron discharge patterns in humans due to the one-to-one spike relation between motoneuron and muscle fiber action potentials, which allows us to make inferences about motor commands. Persistent inward currents (PICs), which provide gain control of motoneuronal output, are facilitated by monoaminergic input from the brainstem. This monoaminergic input is greatly diffuse, but resulting PICs are highly sensitive to inhibitory inputs. Antagonist muscle stretch, and thus Ia input from the antagonist decreases PIC magnitudes in the decerebrate cat. In the present study, we explored whether estimates of PICs are altered with vibratory input to antagonist muscles in humans. MUs of the tibialis anterior (TA), soleus (SOL), and medial gastrocnemius (MG) were discriminated using high-density surface electromyography and convolutive blind source separation. We estimated PICs using the paired MU analysis technique, which quantifies discharge rate hysteresis (ΔF) by comparing the discharge rate of a lower-threshold MU at the onset and offset of a higher-threshold MU. Participants performed isometric plantarflexion and dorsiflexion contractions to a peak of 30% of maximal voluntary contraction, with 10 s ascending and descending phases. In half of the trials, we applied vibration to the antagonist tendon and found that ΔF in agonist MUs decreased in the presence of vibration. These findings suggest that inhibition from the antagonist muscle, most likely Ia reciprocal inhibition, can reduce discharge rate hysteresis. This provides insights about non-invasive methods potentially capable of dampening PICs in hyperexcitable motoneurons, which are manifest in some neurological impairments.

**KEY POINTS:**

- Persistent inward currents in motoneurons amplify synaptic inputs and thus have a major impact on motor unit firing patterns.
- We show that sustained vibration to the antagonist tendon reduces estimates of persistent inward currents (ΔF) of the contracting muscle in both the plantarflexors and dorsiflexors.
- These findings provide evidence for the important role of sensory input in the control of persistent inward currents in the human.
- Reciprocal inhibition may help refine neuromodulatory commands to tailor motor unit activation to diverse movement patterns and specific tasks, and loss of inhibition may exacerbate symptoms of neurological impairment.

## INTRODUCTION

The term ‘reciprocal inhibition’ was first coined by Sir Charles Sherrington in 1897 with his work entitled “On Reciprocal Innervation of Antagonistic Muscles” (1907). Sherrington’s focus on reciprocal inhibition between flexors and extensors would become fundamental to the understanding of central pattern generators and the overall neural control of movement. Since then, many experiments have examined the functional role of Ia reciprocal inhibition during normal behaviour. In fact, disynaptic Ia reciprocal inhibition is arguably the most extensively studied spinal circuit in humans (Pierrot-Deseilligny & Burke, 2005). Ia inhibitory interneurons receive multiple descending inputs and it is generally thought that motor commands activate Ia interneurons in parallel with motoneurons to provide a convenient mechanism for inhibitory control of antagonists during the activation of agonists (Baldissera *et al*., 1981; Pierrot-Deseilligny & Burke, 2005).

Although once believed to be passive integrators of synaptic input, motoneurons are now known to act like amplifier integrators due to their voltage-sensitive dendritic ion channels. Excitatory synaptic input to the motoneuron is both amplified and prolonged because of persistent inward currents (PICs) (Lee & Heckman, 1996, 2000), which are voltage-gated slow-activating L-type Ca and fast-activating persistent Na currents (Schwindt & Crill, 1980; Hounsgaard & Kiehn, 1985, 1989; Powers & Binder, 2000; Hultborn *et al*., 2003; Li & Bennett, 2003). PICs are activated near threshold (slightly below or above) and can amplify synaptic currents by as much as 3-5 fold (Lee & Heckman, 1996, 2000). The level of PIC activation is highly dependent on neuromodulatory drive from the monoaminergic system (i.e. serotonergic and noradrenergic drive) (Lee & Heckman, 1999, 2000; Harvey *et al*., 2006). However, neuromodulatory drive is widely distributed, simultaneously affecting diffuse motor pools throughout a limb (Skagerberg & Björklund, 1985). Although some behaviours could benefit from strong co-activation about a joint, a mechanism must exist to refine the coordinated activation of muscles for tasks such as locomotion and reaching where antagonist co-activation would be a hindrance. In the decerebrate cat, lengthening of the antagonist muscle causes only small increases in Ia reciprocal inhibition but reduces PICs by about 50% (Hyngstrom *et al*., 2007) suggesting that PICs are highly sensitive to input from antagonist Ia afferent input. Therefore, Ia reciprocal inhibition may provide an intriguing mechanism capable of sculpting motor unit (MU) discharge patterns in the presence of strong neuromodulation from monoaminergic tracts (Heckman *et al*., 2008*a*).

The specific role of reciprocal inhibition in the control of motoneuron excitability has been emphasized recently (Heckman *et al*., 2008*a*; Johnson & Heckman, 2010; Johnson *et al*., 2017), and measures of Ia reciprocal inhibition correlate with the magnitude of PICs in humans (Vandenberk & Kalmar, 2014). The capabilities of reciprocal Ia input to dampen the magnitude of PICs in humans, however, has not been extensively tested. Mesquita and colleagues (2022) very recently showed that electrical stimulation to the common peroneal nerve at low frequencies reduces estimates of PICs in the human gastrocnemius, but the inverse was not tested (i.e. tibial nerve stimulation effects on dorsiflexor PICs). Ia reciprocal inhibition has been, however, well described in humans by examining the responses to conditioning stimuli delivered to mixed nerves innervating the antagonist muscle. These responses have been reported as reductions in 1) Hoffmann (H-) reflex amplitude (Crone *et al*., 1990), 2) ongoing EMG activity (Capaday *et al*., 1990), or 3) post-stimulus time histograms (PSTHs) of single MU activity (Ashby & Zilm, 1982; Mao *et al*., 1984; Ashby & Wiens, 1989). Rapid advances in technology have enabled sampling from tens of concurrently active MUs from many muscles, by using high-density surface electromyography (HD-sEMG) array electrodes and blind source separation algorithms (Holobar *et al*., 2009; Negro *et al*., 2016; Del Vecchio *et al*., 2020). Yavuz et al (2018) combined HD-sEMG decomposition with stimulation of the antagonist muscle afferents and both peristimulus frequencygrams (i.e. PSFs) and PSTHs to examine the mutual transmission of Ia reciprocal inhibition between the tibialis anterior (TA) and triceps surae (soleus [SOL]; and, medial gastrocnemius [MG]) MUs. The probability density distributions of reflex activity were fourfold larger in the TA MUs compared to SOL and MG, suggesting an asymmetric distribution of reciprocal inhibition about the ankle joint. Therefore, it seems logical to assume that Ia reciprocal inhibition would be asymmetrical with regards to its role in the control of antagonist PICs in the human lower limb.

In the present manuscript, we tested whether strong activation of antagonist muscle afferents with tendon vibration can alter the MU discharge characteristics in a large population of concurrently active MUs during isometric contractions. We hypothesized that, since tendon vibration causes strong Ia afferent activity, we would cause reciprocal inhibition of the antagonist MUs and reduce estimates of PIC magnitude (i.e. discharge rate hysteresis; ΔF). This would support the hypothesis that Ia reciprocal inhibition plays a specific role in controlling PICs during normal human motor behaviour. In addition, since asymmetry in the magnitude of Ia reciprocal inhibition between the flexors and extensors has been observed previously, we compared the effects of antagonist vibration across opposing muscles acting about the ankle joint. We hypothesized that, based on the recent work of Yavuz and colleagues (2018), the relative effects of antagonist vibration would be greater in the dorsiflexor (i.e. TA) compared to plantarflexor (i.e. MG and SOL) muscles of the ankle.

## METHODS

### Participants and ethical approval

Eleven healthy volunteers (26.8 ± 1.7 years; 175.9 ± 14.4 cm; 73.1 ± 16.8 kg) of both sexes (3F, 8M) participated in the experiment. All participants had no history of cardiovascular, metabolic, or neuromuscular impairment and provided written informed consent prior to partaking in any experimental protocols. The Institutional Review Board of Northwestern University in accordance with the Declaration of Helsinki approved the protocol (STU00202964). A portion of this data (i.e., MU spike trains from the control condition) has been published elsewhere (Beauchamp *et al*., 2022).

### Overview of the main experimental protocol – sustained vibration

Our primary goal was to quantify the discharge characteristics of human MUs during a ramp contraction in two conditions: 1) antagonist muscle vibration, and 2) control. To accomplish this, we had participants first perform two maximal voluntary isometric contractions (MVC) of the plantar- and dorsiflexors interleaved with 2 minutes of rest. For ramp contractions, participants were instructed to produce torque in the intended direction (i.e., plantarflexion or dorsiflexion) by increasing and decreasing at a rate of 3% MVC/s to a peak torque of ∼30% MVC. Prior to beginning the experimental conditions, participants were required to practice each condition at least 5 times to ensure smooth contractions. We then had participants randomly perform six trials of two successive contractions (12 total ramps) under each of the following conditions with at least a minute between trials: 1) plantarflexion, 2) plantarflexion with vibration to the distal TA tendon, 3) dorsiflexion, and 4) dorsiflexion with vibration applied to the Achilles tendon for a total of 48 ramp contractions. The experimental setup and an outline of the conditions is shown in **Figure 1**.

**Figure 1:**
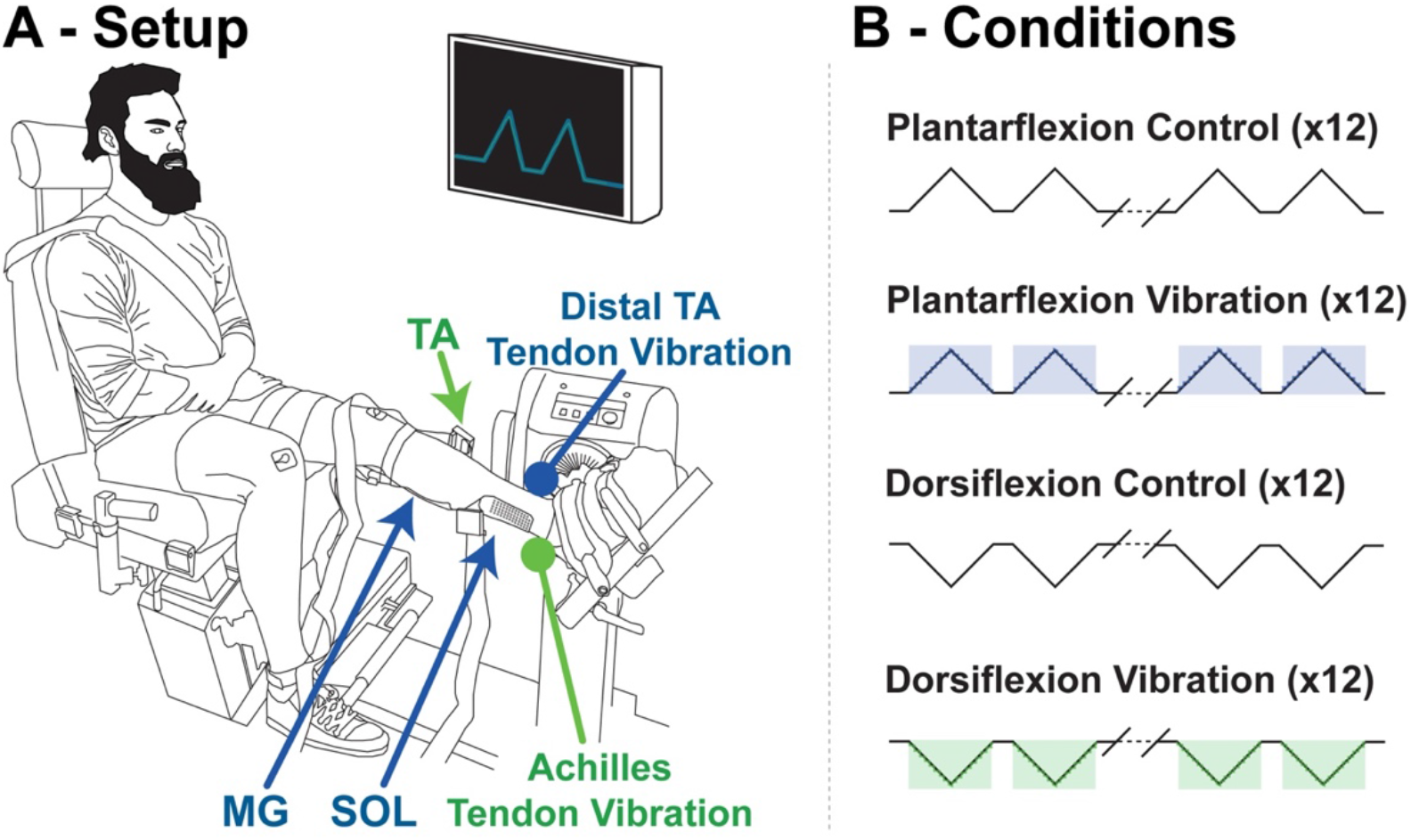
The experimental setup is displayed in A. Note that the muscle recorded from during an intended action is coloured in accordance with the location of vibration used to cause inhibition. Blue corresponds to plantarflexion (i.e. medial gastrocnemius [MG] and soleus [SOL] activation) and distal tibialis anterior [TA] tendon vibration, whereas green corresponds to dorsiflexion (i.e. TA activation) and Achilles tendon vibration. In B, the 4 experimental conditions for the main experimental protocol are shown.

### Overview of the additional experimental protocol – brief vibration

After completion of the ramp contractions, a subset of 5 participants performed ramp and hold contractions in each direction with a 10 s increase, 5 s plateau phase once they reached 30% of MVC, and then 10 s decrease to rest. The purpose of these contractions was to examine the effects of a brief (i.e. 250 ms) period of vibration on MU discharge. First, participants practiced the contractions with and without visual feedback during the last 4 s of the plateau. Once they could maintain torque within ±3% MVC of the target without visual feedback, they performed 4 dorsiflexion trials, in which we randomly applied vibration to the Achilles tendon in 2/4 of the trials. Finally, they performed 4 trials of plantarflexion, in which we randomly applied vibration to the distal TA tendon in 2/4 of the trials. In all cases, participants were told to not intervene if they perceived that the vibration caused them to deviate from the intended target.

### Experimental procedures

#### Torque

Participants were seated comfortably in an isokinetic dynamometer (Biodex System 4; Shirley, NY) with the hips at ∼80 degrees of flexion, left knee at 10 degrees flexion, and the left ankle at 10 degrees of plantarflexion (see **Figure 1**). We secured the left foot to a footplate attached to a Systems 2 Dynanometer (Biodex Medical Systems, Shirley, USA) with the axis of rotation aligned to the center of the ankle joint. Shoulder and thigh straps secured participants to the chair in order to minimize changes in body position. A target line as well as visual torque feedback was provided to the subjects on a 42-inch monitor placed ∼1.5 m in front of the subject at eye level using custom written Matlab software (MATLAB R2020b, The Mathworks Inc., Natick, USA). Torque signals were amplified (150 times), digitized (2048 Hz) and bandpass filtered (10-900) using a 16-bit analog-to-digital converter (Quattrocento, OT Bioelettronica, Turin, IT). Torque signals were recorded with the software OTBioLab+ (Version 1.4, OT Bioelettronica, Turin, IT).

#### High-density surface electromyography (HD-sEMG)

HDsEMG signals were collected from the TA, SOL, and MG using 64 channel electrode grids (GR08MM1305, OT Bioelettronica, Turin, IT) placed on the belly of the muscle and fixed to the skin with adhesive foam (KITAD064, OT Bioelettronica, Turin, IT). Prior to placing the electrode arrays, the skin over the muscles of the lower left leg was shaved, rubbed with abrasive paste, and wiped with isopropyl alcohol. The locations of the muscles were identified through palpation by a clinical exercise physiologist. After ensuring high signal to noise ratios, Hypafix^®^ tape (BSN Medical Inc., Hamburg, DE) was used to minimize movement of the arrays. The array consisted of 64 gold coated electrodes (13 rows x 5 columns) with 1 mm diameter and 8 mm inter-electrode distance (i.e.d.) (GR08MM1305, OT Bioelettronica, Inc., Turin, IT). We securely fixed a band electrode dampened with water around the right ankle, and two Ag/AgCl ground electrodes bilaterally on the right and left patella. HDsEMG signals were acquired with differential amplification (150 times), digitized (2048 Hz) and bandpass filtered (10-900 Hz) using a 16-bit analog-to-digital converter (Quattrocento, OT Bioelettronica, Turin, IT). We visualized and recorded HDsEMG signals with OTBioLab+ (Version 1.4, OT Bioelettronica, Turin, IT) simultaneously with the torque signals.

#### Vibration

A commercially available personal massager (Model 91, Daito-Thrive, Showa-cho, JP) was used to apply vibration to the antagonist tendon of the contracting muscles by manually holding the vibrator perpendicular to the distal TA or Achilles tendon. A 3D-printed sleeve custom-fitted with a load cell (TAS606, HT Sensor Technology Co., Ltd., Shaanxi, CN) made contact (Diameter: 29 mm) with the skin, vibrating at 128 Hz. In all cases, the device was pressed against the skin with 15 ± 2 N of force throughout the contractions. During the main experimental protocol (sustained vibration), vibration commenced and ceased at the onset and offset of the ramp contraction, respectively. During the additional protocol involving brief vibration during ramp and hold contractions, vibration was triggered 2.5 s into the plateau phase of the contraction for a duration of 250 ms.

### Data analysis

#### Torque data

We converted torque data from voltage to torque (Nm) offline and zeroed it to the pre-contraction rest period. Torque signals were low-pass filtered with a third-order, zero-lag Butterworth filter with a cut-off frequency of 10 Hz. All ramp contractions that reached a peak torque between 27 and 33% MVC without substantial deviations from the intended torque target were included in the analysis, whereas those that fell outside of this range or had substantial deviations were excluded (less than 2/48 of contractions per participant).

#### Decomposition of MU spike trains from HDsEMG

Prior to decomposition, surface EMG channels were bandpass filtered at 20–500 Hz (second-order, Butterworth) and then visually inspected. Channels that had substantial artifacts, noise, or saturation of the A/D board were manually removed (usually 2-3 out of 63 bipolar channels). The remaining EMG channels were decomposed into individual MU spike trains using convolutive blind source separation (Holobar *et al*., 2009; Negro *et al*., 2016) and successive sparse deflation improvements (Martinez-Valdes *et al*., 2017). The silhouette threshold for decomposition was set to 0.87 and to ensure we avoided duplicate MUs, we cross-correlated MUs from within the same trial and excluded the decomposed MU with higher covariance from pairs with >30% overlap in spike times. The decomposition accuracy was then improved by manual editing of consecutive spikes by iteratively re-estimating the interval pulse train in order to correct for extraneous deviations in the frequency profile (Boccia *et al*., 2019; Afsharipour *et al*., 2020; Del Vecchio *et al*., 2020; Hassan *et al*., 2021; Martinez-Valdes *et al*., 2020). All the MU spike trains were then visually inspected and only those with a silhouette value that remained > 0.87 were retained for further analysis (Negro *et al*., 2016; Martinez-Valdes *et al*., 2017). In the cat, these MU discharge times have a rate of agreement of ∼90% between MUs decomposed from both the array and fine wire signal (Thompson *et al*., 2018; Kim *et al*., 2020). To ensure we were comparing the same MUs across conditions, we used a normalized cross-correlation between MU action potential profiles across the grid as a measure of similarity, as was introduced by Martinez-Valdes and colleagues (Martinez-Valdes *et al*., 2017). We ran a semi-blind cross-correlation between each control trial and all vibration trials. MU pairs with a normalized cross-correlation higher than 0.8 were considered a matched unit. If multiple matches were found in the same trial (i.e. multiple normalized cross-correlation values >0.8), the algorithm matched the MUs that were cross-correlated with the highest value. This process maximizes the probability of finding the correct MU across conditions and has been validated across days and even weeks (see Martinez-Valdes et al., (2017) and Del Vecchio et al., (2019)).

#### Data analysis

Interspike intervals of each spike train were used to create discharge rate profiles that were filtered and smoothed using support vector regression (for details, see Beauchamp et al., (2022)). The initial (IDR_initial_) and final (IDR_final_) discharge rates were quantified as the value from the smoothed discharge rate at the time instance of the first and last spikes in the MU spike train, whereas the peak (IDR_peak_) discharge rate was the highest value identified in the smoothed discharge rate profile. The torque at recruitment and de-recruitment were quantified as the instantaneous torque at the instance of the first and last spikes in the MU spike train.

We used the paired MU analysis technique to estimate the effects of PICs on motoneuron discharge patterns by measuring the discharge hysteresis of a higher threshold (test) MU with respect to the discharge rate of a lower threshold (reporter) unit (Gorassini *et al*., 2002). More specifically, ΔF was quantified as the difference in the discharge rate of the reporter unit at test unit recruitment and de-recruitment. To allow full activation of the PIC in the reporter unit, we excluded any pairs with recruitment time differences <1 s (Bennett *et al*., 2001; Powers *et al*., 2008; Hassan *et al*., 2020). Furthermore, to ensure MU pairs received common synaptic drive, we only included test unit-reporter unit pairs with rate-rate correlations of r^2^ > 0.7 (Gorassini *et al*., 2004; Udina *et al*., 2010; Stephenson & Maluf, 2011; Vandenberk & Kalmar, 2014). Finally, we excluded test unit-reporter unit pairs in which the reporter unit discharge range (DRmax – DRmin) was < 0.5 pps while the test unit was active (Stephenson & Maluf, 2011). When utilizing decomposition of HDsEMG to identify MUs, each test unit can have tens of usable test unit-reporter unit pairs that can be used to calculate ΔF. To reduce the number of ΔF estimates obtained from each unit, while also reducing the variability across the pool of identified MUs, we chose to take the mean of all ΔF values obtained from each test unit that satisfied our inclusion criteria (Khurram *et al*., 2021; Hassan *et al*., 2021). This value will henceforth be referred to as simply ΔF. We provide a general schematic of the process we used to obtain this measure of ΔF in ***Figure 2***.

**Figure 2:**
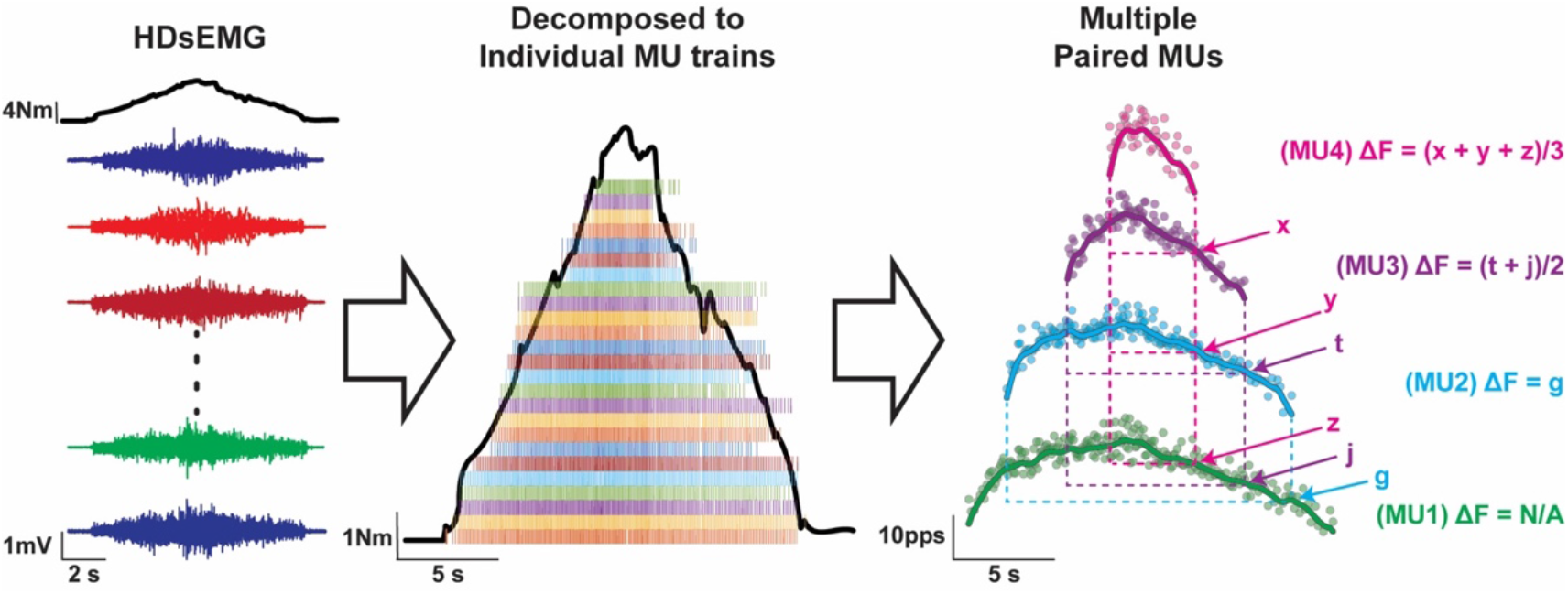
The process of decomposition from multiple channels of surface EMG (left) to individual motor unit spike times (middle), to quantifying ΔF in many motor units (right). In this reduced example, there are three test units that can be examined. The ΔF for MU4 (pink) is derived from the average of ΔF between reporter units MU1 (green), MU2 (blue), and MU3 (purple). The ΔF for MU3 (purple) is derived from the average of ΔF between reporter units MU1 (green) and MU2 (blue). The ΔF for MU2 (blue) is derived from the ΔF with reporter unit MU1 (green) only.

To quantify the relative inhibitory effects of vibration applied briefly to the antagonist tendons during plantarflexion and dorsiflexion contractions (i.e., additional experiment – brief vibration), we compared the mean smoothed discharge rate during a 100 ms window 125-25 ms prior to and 25-125 ms following the onset of vibration. This allowed us to determine the effects of brief vibration on the average behaviour of the MU pool within a subject.

### Statistical Analysis

All statistical procedures were performed using R (Version 3.3.3, The R Foundation, Vienna, Austria) and R Studio (Version 4.0.2, RStudio Inc., Boston, USA). In the main experiment, we detail the effects of sustained antagonist tendon vibration on MU discharge characteristics between control and antagonist vibration ramp contractions by comparing only those MUs that could be reliably matched between vibration and control contractions. We chose to use linear mixed effects models to avoid the assumption that the subject, trial, contraction number within a trial, and individual MU within a participant did not influence the outcome variables. In doing so, we take into consideration all our data points (i.e., within-subject and trial variability) rather than averaging across them and basing our analysis on the mean within an individual trial or subject (Addelman, 1970; Hopkins, 1982; Moerbeek *et al*., 2003; Boisgontier & Cheval, 2016; Giboin *et al*., 2020; Hassan *et al*., 2021; Khurram *et al*., 2022). Linear mixed effects model analysis was performed in R using the lme4 package (Bates *et al*., 2015) and significance was calculated using the lmerTest package (Kuznetsova *et al*., 2017), which applies Satterthwaite’s method to estimate degrees of freedom and generate p-values for mixed models. More specifically, we employed linear mixed effects models of IDR_initial_, IDR_peak_, IDR_final_, and ΔF to examine their relationship with antagonist vibration across the three sampled muscles in units that could be matched across conditions. As fixed effects, we included condition (antagonist vibration and control), muscle (SOL, MG, and TA), and their interaction terms. As random effects, we included random intercepts for subject, trial number, ramp number, and individual MU identifier. We also analyzed whether the condition and muscle can predict the changes in the torque at MU recruitment and de-recruitment across conditions. In this case, we included muscle, condition, and their interaction as fixed effects, and we included subject, trial, ramp number, and unit identifier as random effects.

For all mixed models, we confirmed that the assumptions of linearity, homoscedasticity and normality of residuals were valid for all data. We obtained p-values for individual models by likelihood ratio tests of the full model with the fixed effects in question against models without those effects. To provide a standardized effect of antagonist vibration-induced differences for each muscle, we computed Hedge’s G effect sizes from the estimated marginal means (Searle *et al*., 1980) that were obtained from our mixed effects models using the emmeans package (Lenth *et al*., 2022).

#### Supplementary experiment – brief vibration

To determine the effects of brief antagonist tendon vibration on MU discharge, we compared the mean discharge rate of all concurrently active MUs immediately before, and during 250 ms of vibration to the antagonist tendon. To do this, we used a linear mixed effects model to examine the relationships between discharge rate and antagonist tendon vibration across the three muscles that we sampled. We employed time point (pre or during vibration), muscle (TA, MG or SOL), and their interaction as fixed effects. As random effects, we included random intercepts for subject, trial, and MU. Hedge’s G effect sizes were calculated to provide a standardized effect of antagonist vibration-induced differences for each muscle. All values reported in text and figures are means ± standard deviation.

## RESULTS

### Motor unit decomposition

A total of 1131 and 1147 MU spike trains were identified across all participants in the TA muscle during the control and vibration trials, respectively, of which 470 pairs were matched across conditions. A total of 1511 and 1125 MU spike trains were identified across all participants in the MG muscle during the control and vibration trials, respectively, of which 318 pairs were matched across conditions. A total of 833 and 768 MU spike trains were identified across all participants in the SOL muscle during the control and vibration trials, respectively, of which 202 pairs were matched across conditions.

### Peak torque achieved across submaximal triangular conditions

Although participants provided anecdotal reports of increased difficulty when producing torque in the antagonist vibration conditions, peak plantarflexion torque achieved during the submaximal ramps was similar between control (28.3 ± 1.12% MVC) and vibration (28.2 ± 0.89% MVC) trials. Likewise, peak dorsiflexion torque achieved during the submaximal ramps was similar between control (29.5 ± 0.96% MVC) and vibration (30 ± 1.32% MVC) trials.

### Antagonist tendon vibration dampens ΔF irrespective of the tested muscle

A summary of the effects of antagonist tendon vibration on all variables of interest is provided in **Table 1**. Muscle (χ^2^(1) = 175.52, p < 0.0001) and condition (χ^2^(1) = 14.289, p = 0.00016) were significant predictors of ΔF when we examined MUs matched across conditions, however their interaction was not (χ^2^(2) = 3.913, p = 0.1414). SOL (2.87 ± 0.257 pps) ΔF values were lower (p < 0.00012) than those of MG and TA, and MG (3.55 ± 0.25 pps) ΔF values were lower (p < 0.0001) than those of TA (4.81 ± 0.25. pps). Regardless of muscle, ΔF was lower during antagonist vibration by approximately 0.54 ± 0.09 pps (see **Figure 3**). In **Figure 4** we provide example trials showing that ΔF values of individual matched units across conditions decrease with antagonist muscle tendon vibration for the TA.

**Table 1:**
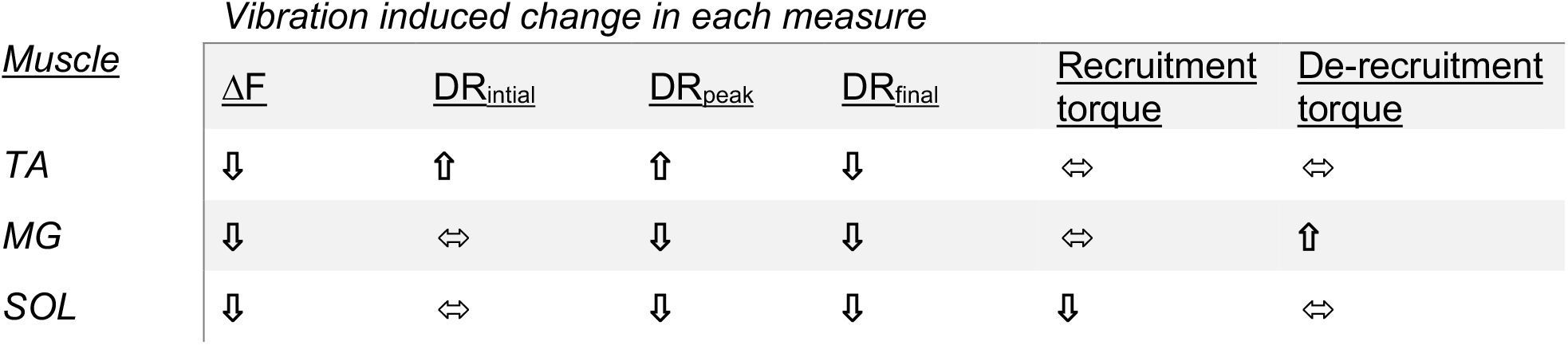
Summary of antagonist tendon vibration induced changes in motor unit behaviour during ramp conditions. Arrows indicate either a significant increase (⇑), significant decrease (⇓), or no change (⇔). TA = tibialis anterior; MG = medial gastrocnemius; SOL = soleus; ΔF = discharge rate hysteresis; DR = discharge rate.

| <u>Muscle</u> | <u>Vibration induced change in each measure</u> |  |  |  |  |  |
| --- | --- | --- | --- | --- | --- | --- |
|  | <u>$\Delta F$</u> | <u>$DR_{initial}$</u> | <u>$DR_{peak}$</u> | <u>$DR_{final}$</u> | <u>Recruitment torque</u> | <u>De-recruitment torque</u> |
| TA | ↓ | ↑ | ↑ | ↓ | ↔ | ↔ |
| MG | ↓ | ↔ | ↓ | ↓ | ↔ | ↑ |
| SOL | ↓ | ↔ | ↓ | ↓ | ↓ | ↔ |

**Figure 3:**
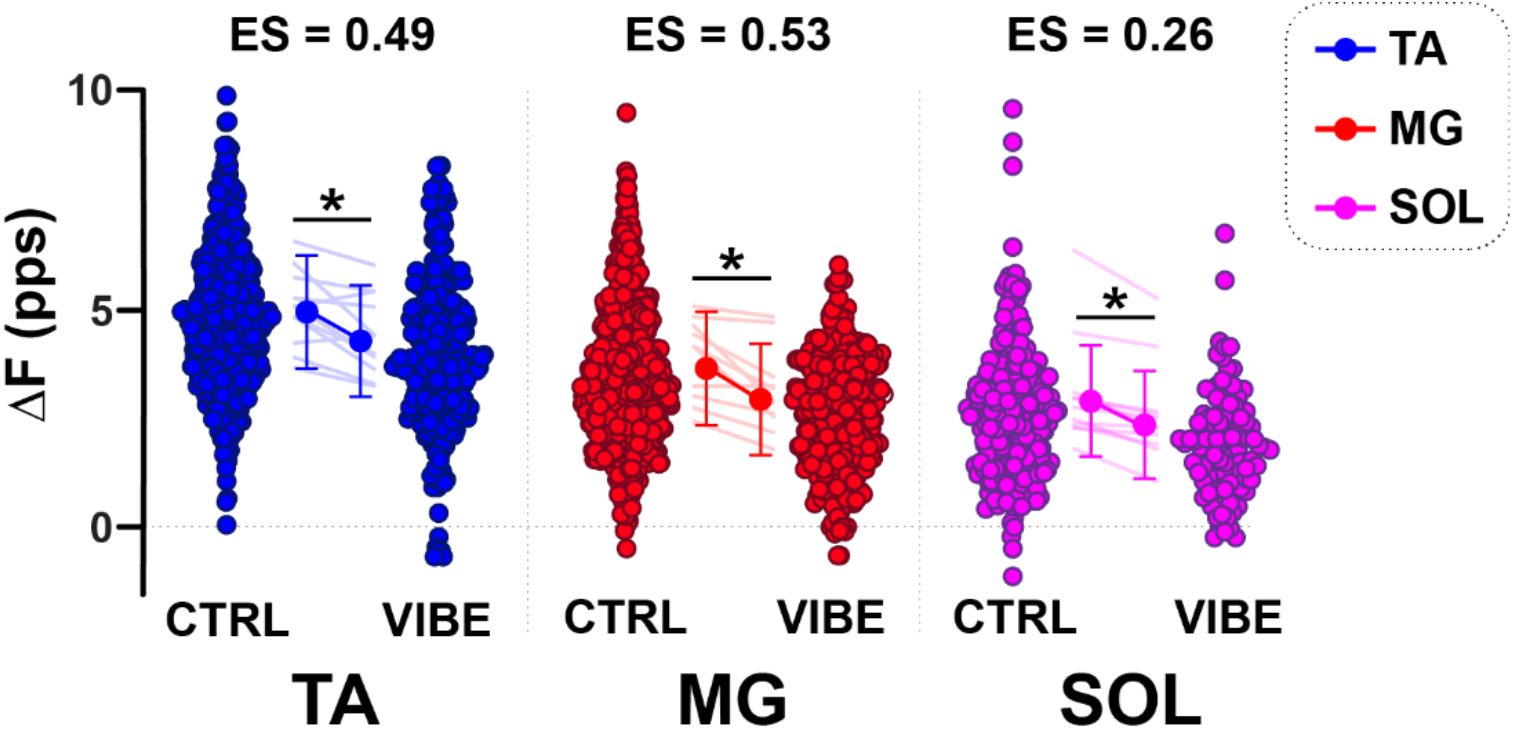
Test unit mean discharge rate hysteresis (ΔF) values for the tibialis anterior (TA), medial gastrocnemius (MG) and soleus (SOL) motor units during ramps contractions in the control (CTRL) and antagonist vibration (VIBE) conditions. Each point in the distributions is from one unit that was matched between conditions. Points in the center represent the model estimates from the linear mixed effects model and the error bars are the corresponding 95% confidence intervals. These model estimates are superimposed on the mean change between conditions for all units within a participant (transparent lines). Asterisks indicate significant difference between conditions (p < 0.05).

**Figure 4:**
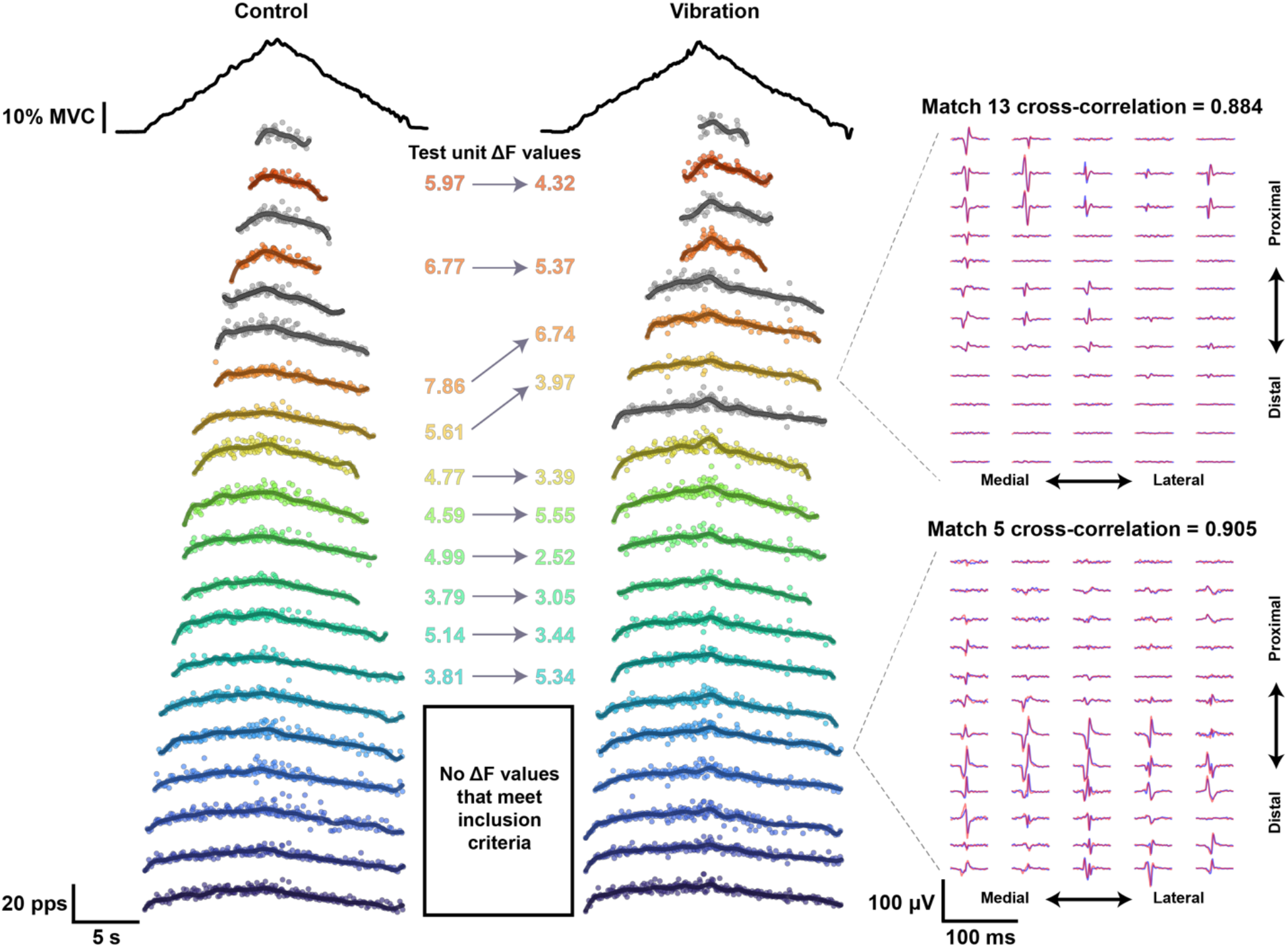
An example of matched unit analysis between conditions from one trial in each condition. In this case, 20 out of 28 motor units are shown in the control trial and 20 out of 29 units are shown in the vibration trial. For each condition, the torque traces are shown at the top. Below each torque trace, matched units are color-coded between conditions and organized from highest- to lowest-threshold (warmer to cooler colors, respectively). An acceptable match was not found for motor unit firing patterns shown in grey. Each point represents the instantaneous discharge rate, with the overlaid lines representing the smoothed discharge rate computed using support vector regression. Discharge rate hysteresis (ΔF) values for each matched test unit is shown in between the trials and individual matched units are connected by an arrow. In this case, 16 of 20 of the units shown were matched using a two-dimensional cross-correlation (Martinez-Valdes et al., 2017). This process is shown to the right for two different matched units. The individual motor unit action potentials (MUAPs) obtained from spike triggered averages at the instances of motor unit discharge are arranged across the orientation of the high-density surface electromyographic array. Control MUAPs are shown in blue, whereas MUAPs from the vibration trial are shown in red.

### Effects of antagonist vibration on initial, peak, and final discharge rates

Figure 5 displays ensemble averages (Beauchamp *et al*., 2022) of the smoothed discharge patterns from MUs in the entire dataset and illustrates the trends across all decomposed MUs. Linear mixed effects models indicated antagonist vibration-induced changes in ***initial discharge rates*** were predicted by a significant interaction between condition and muscle (χ^2^(5) = 259.01, p < 0.0001). ***Initial discharge rates*** were higher during antagonist vibration in the TA (control = 5.61 ± 0.205 pps; vibration = 5.95 ± 0.209 pps; p = 0.0128) but did not differ between conditions for SOL (p = 0.1811) or MG (p = 0.9952). Across conditions, ***initial discharge rates*** were different for each muscle (TA = 5.76 ± 0.201 pps; MG = 4.64 ± 0.201 pps; SOL = 4.06 ± 0.205 pps; all p < 0.0001).

**Figure 5:**
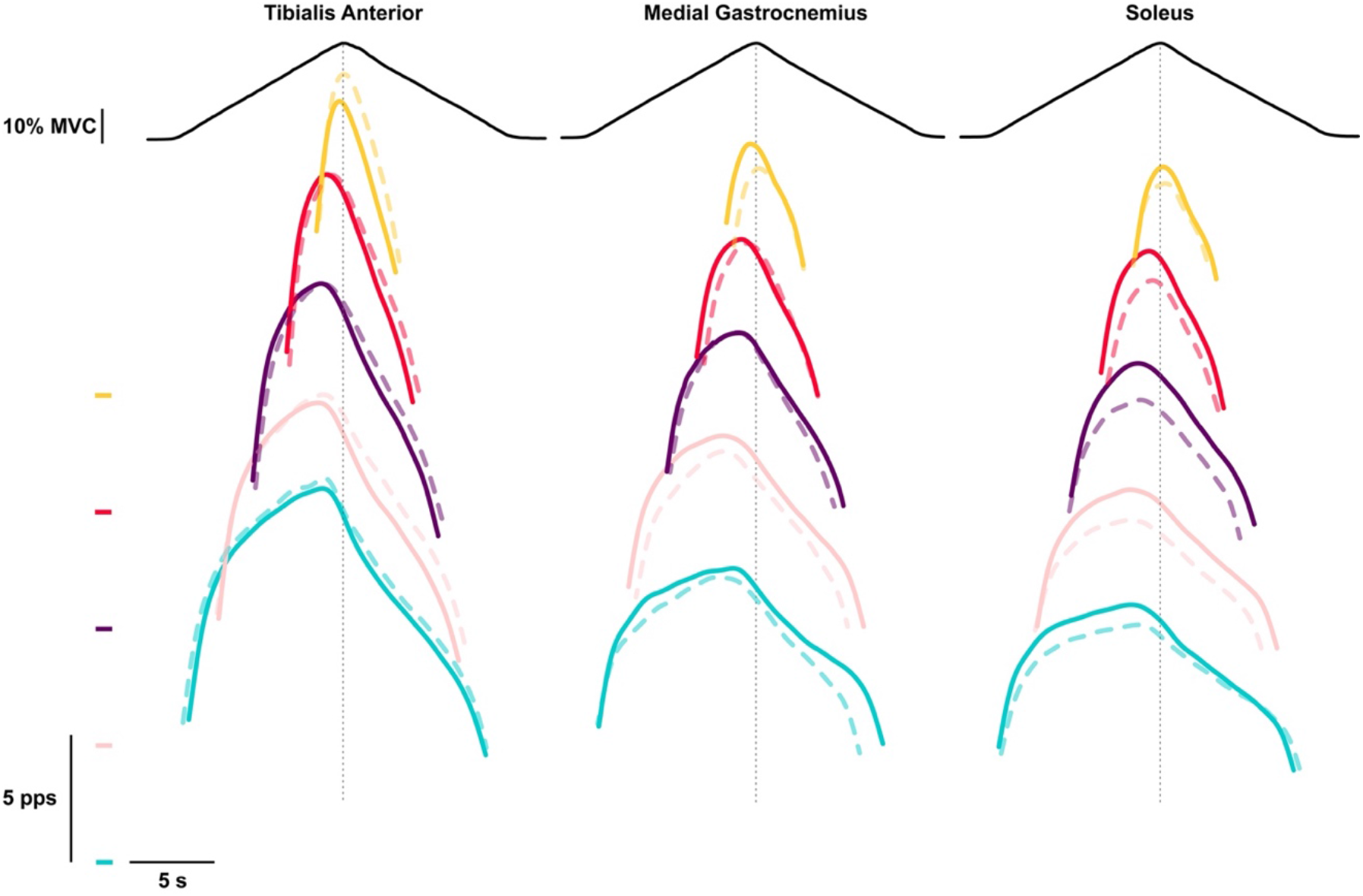
Averaged torque and discharge rates obtained from all motor units during the control (solid lines) and vibration (dashed lines) conditions. Ensembles were created using methods that are previously described (Beauchamp et al., 2022). Motor units are stratified by recruitment thresholds across participants; 0-6% (teal), 6-12% (pink), 12-18% (purple), 18-24% (red), and >24% (yellow) of MVC. The zero for instantaneous discharge of each ensemble is indicated to the left with corresponding coloured lines. The vertical dashed lines indicate the peak of torque. From these plots, one can appreciate differences in the mean behaviour of units across muscles and conditions.

Antagonist vibration-induced changes in ***peak discharge rates*** were predicted by the interaction between condition and muscle (χ^2^(5) = 388.67, p < 0.0001). ***Peak discharge rates*** of the TA increased slightly with antagonist vibration (control = 16.3 ± 0.606 pps; vibration = 17.2 ± 0.611 pps; p = 0.01), whereas antagonist vibration caused a decrease in ***peak discharge rates*** of SOL (control = 11.8 ± 0.61 pps; vibration = 10.7 ± 0.61 pps; p = 0.0008) but had no significant effects on MG MUs (control = 12.7 ± 0.6 pps; vibration = 12 ± 0.61 pps; p = 0.149).

Antagonist vibration slightly reduced ***final discharge rates*** in MUs of all three muscles (χ^2^(1) = 8.5398, p = 0.0035) but the interaction between condition and muscle was not significant (χ^2^(2) = 2.7393, p = 0.2542). Across muscles, ***final discharge rates*** were reduced (p = 0.0033) during the vibration (6.09 ± 0.234 pps) compared to control (6.41 ± 0.231 pps) condition. Across conditions, ***final discharge rates*** were predicted by the muscle (χ^2^(2) = 233.45, p < 0.0001) and different between muscles (TA = 7.53 ± 0.235 pps; MG = 6.06 ± 0.235 pps; SOL = 5.16 ± 0.24 pps; all p < 0.0001).

### Effects of antagonist vibration on the torque at recruitment and de-recruitment

Linear mixed effects analysis indicated that ***recruitment*** torque was predicted by the interaction between condition and muscle (χ^2^(5) = 95.624, p < 0.0001). With antagonist vibration, the torque at ***recruitment*** was reduced for SOL (control = 17.4 ± 0.65% MVC; vibration = 15.7 ± 0.7% MVC; p = 0.0048) but there was no difference between conditions for MUs in the MG (control = 19.8 ± 0.62% MVC; vibration = 19.9 ± 0.65% MVC; p = 0.999) or TA (control = 21 ± 0.63% MVC; vibration = 20.5 ± 0.67% MVC; p = 0.876).

Similarly, ***de-recruitment*** torque was predicted by the interaction between condition and muscle (χ^2^(5) = 68.475, p < 0.0001). With antagonist vibration, the torque at ***de-recruitment*** was increased for MG (control = 6.9 ± 0.5% MVC; vibration = 8.8 ± 0.54% MVC; p = 0.0009) but there was no difference between conditions for MUs in the SOL (control = 4.7 ± 0.54% MVC; vibration = 5.59 ± 0.61% MVC; p = 0.5299) or TA (control = 6.34 ± 0.51% MVC; vibration = 5.22 ± 0.57% MVC; p = 0.2288).

### Supplementary experiment – the inhibitory effect of brief antagonist tendon vibration on discharge rates is greater in the TA than SOL and MG

After noting that sustained vibration was symmetrical between opposing muscles about the ankle, we sought to determine whether the inhibitory effects of brief antagonist vibration to ongoing MU activity in the plantarflexors and dorsiflexors was symmetrical. In doing so, we provided brief vibration to the Achilles tendon during dorsiflexion and distal TA tendon vibration during plantarflexion. Our linear mixed effects model showed that the interaction between muscle and time point (prior to vs during vibration) was a significant predictor of discharge rate (χ^2^(5) = 443.3, p < 0.0001). Discharge rates were reduced during Achilles tendon vibration for TA (before = 13.56 ± 0.99 pps; during = 11.87 ± 1.0 pps; p < 0.0001) but there were no changes in MU discharge rates of the MG (before = 9.79 ± 0.99 pps; during = 9.29 ± 0.93 pps; p = 0.177) or SOL (before = 9.32 ± 1.02 pps; during = 9.02 ± 1.0 pps; p = 0.9115) with distal TA tendon vibration.

## DISCUSSION

The present study aimed to investigate the effects of antagonist tendon vibration on MU firing patterns during submaximal isometric ramp contractions in the human lower limb. In agreement with recent literature (Mesquita *et al*., 2022), we found that estimates of persistent inward currents (PICs; ΔF) were significantly lower in the presence of sensory input from the antagonist muscle. Furthermore, and perhaps most novel, we found symmetry across muscles in in terms of reduced estimates of PICs. That is, vibration to the distal TA and Achilles tendons similarly reduced estimates of PICs in the triceps surae (i.e., SOL and MG) and TA, respectively. This contrasts with the inconsistent effects of antagonist tendon vibration on initial, peak and final discharge rates, and on recruitment and de-recruitment thresholds that we observed across muscles (see Table 1). Brief vibration applied to the antagonist tendon did, however, have asymmetrical effects on MU firing rates, like that of electrical nerve stimulation-induced reciprocal inhibition (Yavuz *et al*., 2018). These findings suggest that, despite the inconsistent and asymmetrical effects on MU recruitment and firing patterns across muscles, antagonist tendon vibration consistently reduces estimates of PICs, which may have implications for clinical scenarios where dampening of PICs in hyperexcitable motoneurons may have a therapeutic role.

### Methodological considerations

Vibration is a crude method of inducing Ia input and there are several other types of sensory input resulting from vibration applied to the antagonist tendon. For instance, vibration excites mechanoreceptors both locally and non-locally, and cutaneous receptors have shown powerful effects on MU recruitment (Garnett & Stephens, 1981). Vibration applied to the antagonist tendon could also provide Ia input to the agonist and heteronomous muscle spindles, which could affect MU excitability. For instance, indirect vibration causes reductions in Hoffmann reflex amplitudes and increased amplitudes of the inhibitory middle latency cutaneous reflex amplitudes (Barss *et al*., 2021). Despite these concerns, we found asymmetrical effects of brief antagonist tendon vibration on MU discharge rates between the TA and triceps surae, like the asymmetrical effects of peripheral nerve stimulation to induce Ia reciprocal inhibition (Yavuz *et al*., 2018), indicating that we were probably activating Ia afferents like stimulation of peripheral nerves innervating muscle spindles of the antagonist muscle.

The task performance may have diminished the magnitude of the inhibitory effects of antagonist tendon vibration on MU discharge and estimates of PICs. Since the participants were required to reach the same absolute torque in each condition, it is possible and likely that they provided additional voluntary drive during the antagonist tendon vibration conditions. This is evidenced by the anecdotal reports of the increased difficulty of reaching 30% of MVC during the antagonist tendon vibration conditions. This additional voluntary drive (i.e., excitatory synaptic input) may have overcome the inhibitory actions of antagonist tendon vibration (i.e., Ia reciprocal inhibitory input) to motoneurons of the contracting muscles. Indeed, this may have resulted in recruitment of additional MUs to achieve the same torque or, in the case of the TA, increased discharge rates of higher threshold units that was mediated by additional excitatory synaptic input to compensate for the reduced contribution of PICs to force output.

### Consistent and symmetrical dampening of PIC magnitude with antagonist tendon vibration between muscles

Although incorrect, we hypothesized that the dampening effects of antagonist tendon vibration on estimates of PICs would be greater from the Achilles tendon to the MUs of the TA than the reverse (i.e., TA tendon onto triceps surae MUs). The rationale for this hypothesis was two-fold: First, 1) vibration of the muscle or tendon can produce inhibitory postsynaptic potentials onto antagonist motoneurons through Ia inhibitory interneurons (Heckman & Binder, 1991); 2) PICs are highly sensitive to reciprocal inhibition (Hyngstrom *et al*., 2007); and 3) estimates of PICs (i.e., ΔF) are larger in the TA than the SOL (Kim *et al*., 2020). This would suggest that the same inhibitory input provided to the MUs with higher PIC magnitudes would be affected to a greater extent than MUs with lower PIC magnitudes. Since antagonist tendon vibration affected the estimates of PIC magnitude of muscle groups similarly, it is possible that the ΔF value does not accurately depict the absolute magnitude of PICs, and not widely suitable for comparison across muscles. It is possible that the ΔF value may require normalisation (Oya *et al*., 2009) or that other indices of PICs, such as post-acceleration discharge rate attenuation or self-sustained persistent discharge, be used in concert with ΔF to provide a better approximation of the relative contribution of PICs to motoneuron discharge across muscles.

Second, reciprocal inhibition from the triceps surae to TA MUs has been shown to be much stronger than from the TA muscle to triceps surae MUs using short pulses of electrical stimulation delivered to peripheral nerves (Yavuz *et al*., 2018). The functional implications of this reciprocal interaction is believed to relate to neuromechanical adjustments of joint torques about the ankle during postural control and locomotion. Reflexes play a major role in the neuromechanical adjustments required to maintain standing and ensure smooth locomotor behaviour in humans (Zehr & Stein, 1999; Zehr *et al*., 2016; Pearcey & Zehr, 2019, 2020). Although PICs are likely important for both postural control and locomotion (Heckman *et al*., 2008*b*), adjustments in ongoing behaviour are probably not as dependent on PICs. During maintenance of sustained and voluntary torques however, PICs are likely to contribute to a greater degree (Heckman *et al*., 2008*b*). The vibration in the current experiment probably resulted in hyperpolarization of antagonist motoneurons because of inhibitory postsynaptic potentials via Ia inhibitory interneurons (Heckman & Binder, 1991) and this sustained inhibition likely dampened the contributions of PICs to sustained motoneuron discharge equally to all muscles, rather than having asymmetrical effects of brief Ia reciprocal inhibition, like the brief adjustments required during postural and locomotor behaviours. Indeed, our brief vibration results indicated that there was asymmetry in the brief effects of antagonist vibration on motoneuron discharge. TA MUs decreased discharge rate during brief vibration, whereas triceps surae MUs did not. This suggests that the asymmetry of previous results was not due to the method of providing inhibition (i.e. vibration vs. electrical stimulation) but rather the direct effects of inhibition on PIC contributions to motoneuron discharge versus brief effects of Ia reciprocal inhibition on motoneuron discharge.

These considerations beg the question: why does sustained inhibitory input (i.e., vibration of the antagonist muscle) have symmetrical effects of on estimates of PICs, but brief inhibitory inputs (i.e., brief electrical and/or vibratory input to the antagonist) have asymmetrical effects on adjustments in MU discharge? We suggest that the most reasonable explanation is that PICs in triceps surae muscles (i.e., SOL and MG in this study) may have lower voltage thresholds than their dorsiflexor (i.e., TA) counterparts. The PICs of the triceps surae MUs, therefore, depolarize the dendrites to a lesser degree and decrease the impact of transient inhibition from TA and/or extensor digitorum longus (EDL). The prolonged vibration, however, reduces the overall amplitude of the PIC and depolarizes its voltage threshold. This is manifest as a reduced discharge rate hysteresis, as observed in the MUs. This explanation is necessarily speculative, however, and requires further investigation.

### Potential for clinical implications

Current approaches for controlling spasms in people with spinal cord injuries is primarily limited to pharmaceutical approaches, but many individuals do not enjoy taking large quantities of medications due to their lack of effectiveness and negative side effects (McKay *et al*., 2018). The work of DeForest and colleagues (2020) promoted antagonist vibration as a targeted method for controlling spasms in people with spinal cord injuries. They showed that vibration reduced the amplitude of the late component of cutaneous reflexes in antagonist, but not agonist, muscles in people with chronic spinal cord injury. This late component of the cutaneous reflex is mediated almost entirely by Ca^2+^ PICs (Li *et al*., 2004; Murray *et al*., 2010), leading DeForest and colleagues (2020) to suggest that vibration to a muscle may affect PICs in the antagonist muscle. Indeed, our data corroborates this suggestion, and suggests that antagonist tendon vibration may have clinical relevance for the control of spasms. Although we studied the effects of vibration in neurologically intact humans, it remains possible that targeted vibration of antagonist muscles in people with other types of neurological impairment (i.e., hemispheric stroke, cerebral palsy, multiple sclerosis, etc.) may be effective for dampening hyperexcitable motoneuron activity.

## CONCLUSION

The present study examined the relative inhibitory effects of antagonist tendon vibration on the discharge patterns of MUs from the ankle dorsi- and plantarflexors during isometric ramp contractions. Unlike previous reports of transient Ia reciprocal inhibition that show asymmetry of effects on muscles about the ankle (Yavuz *et al*., 2018), we show antagonist tendon vibration caused a significant reduction in estimates of PIC magnitude similarly across muscles. Additional, but inconsistent, effects of antagonist tendon vibration were observed with regards to MU recruitment and discharge rates across muscles. These effects likely occurred because participants were required to compensate for the reduced contribution of PICs to motoneuron discharge. Taken together, the results from the present study suggest that antagonist tendon vibration may prove useful as a non-invasive method for mitigating PIC-induced activity of motoneurons with excessive excitability. This approach may be particularly beneficial for counteracting spastic activity, which is manifest in a variety of neurological impairments or injuries.

## Acknowledgements

The authors would like to thank Ahalya Mandana for technical assistance with the design of the apparatus used in the study.

## Data availability statement

The data that support the findings of this study are available on request from the corresponding author.

## Author contributions

G.E.P.P., O.U.K., and C.J.H. conceptualized and designed the research; G.E.P.P., O.U.K., and J.A.B. performed the experiments; G.E.P.P., O.U.K., J.A.B., and F.N. analysed the data; G.E.P.P., O.U.K., J.A.B., and C.J.H. interpreted the results of experiments; G.E.P.P., O.U.K., and J.A.B. prepared the figures; G.E.P.P. drafted the manuscript; G.E.P.P., O.U.K., J.A.B., F.N., and C.J.H. revised and approved the final version of the manuscript.

